# Scale-resolved correlation functions unify spatial omics across platforms and resolutions

**DOI:** 10.64898/2026.10.09.757915

**Authors:** Mykhailo Ilin, Yerik Singh, Simon W. Mages

## Abstract

Spatial omics platforms populate a multidimensional parameter space and differ in resolution, panel size, sensitivity, and modality. This diversity has led to platform-specific analyses whose results are rarely comparable. Here we present SCOTE (Spatial COvariance TEnsors), a computational framework that quantifies spatial relationships of all feature pairs across all length scales, for molecule coordinates and binned counts alike. Measuring the same quantity, with statistical errors, across platforms, samples, and patients, SCOTE separates measurement effects from biology, with platform artifacts appearing as identifiable, scale-localized features. SCOTE’s molecule-level functions resolved a knockout effect in mouse spleen that cell-level analyses missed, and could show that tissue area and replicates, not molecule depth, set the statistical error. To match the growth of spatial omics data sets, SCOTE runs efficiently on CPUs and GPUs, and processes a billion-molecule whole-transcriptome section at all length scales, with uncertainties, in 26 minutes on a single GPU.

## 1 Introduction

Spatial omics now measures gene expression in intact tissue on many platforms [1]. Imaging platforms such as MERFISH/MERSCOPE, Xenium/Atera, CosMx, and AVITI24 Cytos report the coordinates of individual molecules, for panels up to the whole transcriptome [2–7]; sequencing-based platforms report whole-transcriptome counts per barcoded spot, bead, bin, or spatially localized nucleus [8–13]. Tissue organization, in turn, spans length scales from the sub-micrometer arrangement of transcripts inside a cell to millimeter-sized compartments of an organ [14, 15]. Most analyses nevertheless operate at one scale on a cell-level representation: molecules are first assigned to segmented cells [16, 17], cells are clustered into types, and spatial statistics are computed on cell centroids [18–21], or genes are tested one at a time for spatial variability [22, 23]. Segmentation is the weakest step of this chain: transcript spillover between neighboring cells is widespread on current imaging platforms [24–26], which has motivated segmentation-free analyses of molecules and bins [17, 27–29]. Results obtained on different platforms or with different preprocessing are, moreover, hard to compare quantitatively [30–34].

Pair-correlation statistics are the classical tool for spatial point patterns [35–37] and have been adapted to spatial omics as neighborhood-enrichment and co-occurrence scores of cell types or annotation fractions [18, 19, 29, 38, 39], as Ripley’s *K*, pair-correlation, and mark-correlation functions of cell types with parametric or functional summaries [40–42], and as cross-pair-correlation functions for multiplexed imaging [43]. Most of these methods share three limitations. They operate on segmented and typed cells, so they cannot resolve scales below the cell diameter and inherit segmentation errors. They scale with the square of the number of objects, which restricts them to cells or to subsamples of molecules. Also, they provide no common normalization that would make a number measured on one platform comparable with the same number measured on another and, apart from bootstrap bands for one summary function at a time in spatstat [37, 44] and tests of *K*- or *g*-functions across samples [42, 45–47], no error model attached to every value of every feature pair.

Here we treat every platform’s output as a sample of the same underlying molecular density and measure its two-point correlation function *g*_*ab*_ (*r*): how often molecules of features *a* and *b* occur at distance *r*, relative to the expectation for the observed tissue geometry. Because this function is defined identically for molecule coordinates and for binned counts, we compute it for all feature pairs from the platform’s resolution limit to the size of the section. Three design choices make this practical. A random-label reference [36, 48–51] removes tissue shape, holes, and density gradients without a model; a log-hierarchical lattice turns the pair counts of all feature pairs in a distance shell into one sparse matrix product; and a block bootstrap over tiles [44, 52–55], evaluated in closed form, attaches an uncertainty to every value. We implement this in SCOTE (Spatial COvariance TEnsors), an open-source Python package.

We show that two-point patterns agree more between platforms than between patients, that molecule-level functions resolve a knockout effect that cell-level analyses miss, that tissue area and replicates set the statistical error, that platform artifacts appear as identifiable, scale-localized features, and that a billion-molecule whole-transcriptome section takes 26 minutes on one GPU.

## 2 Results

### 2.1 Molecule-level two-point functions from nanometers to millimeters in one pass

For two features *a* and *b* (genes, or derived “operators” such as cell-type marker sets) we count pairs of molecules, or of counts in bins, at distance *r* and divide by the count expected if the labels were shuffled among the observed positions (Fig. 1; Methods). A value *g*_*ab*_ (*r*) = 1 therefore means “as often as expected for the actual geometry of this section”, and log *g*_*ab*_ (*r*) is the log fold enrichment of cooccurrence at distance *r*: positive for features found together, negative for features that avoid each other. By default, below 4 µm, molecule pairs are enumerated directly; above, molecules are aggregated on a lattice whose pitch doubles from level to level. The pair counts of all feature pairs in one distance shell are a single sparse matrix product, so all pairs at all scales are computed in one pass (Methods).

**Figure 1.**
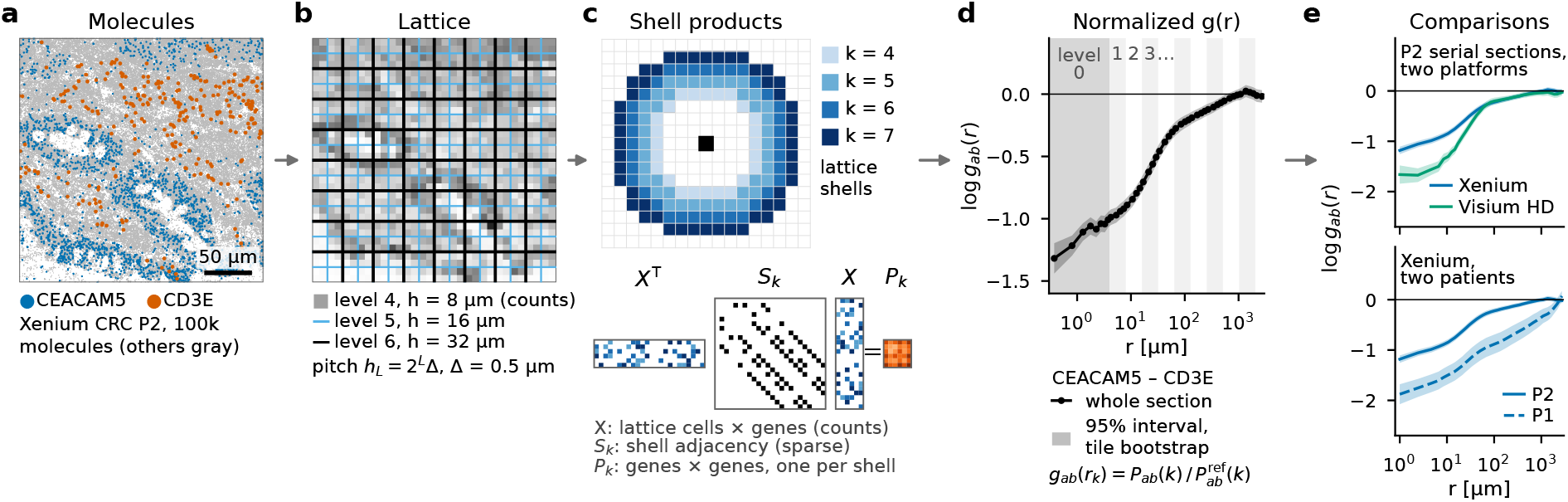
From molecules to normalized two-point functions with errors. **a**, Molecules in a 256 µm crop of Xenium CRC section P2; CEACAM5 (tumor) and CD3E (T cells) colored. **b**, The crop on the log-hierarchical lattice: counts per cell at level 4, cells of levels 5 and 6; the pitch *h*_*L*_ = 2^*L*^Δ (Δ = 0.5 µm) doubles per level. **c**, The four distance shells of one lattice level around a lattice cell, and the shell product *P*_*k*_ = *X*^⊤^ (*S*_*k*_ *X*) of the count matrix *X* with the sparse shell adjacency *S*_*k*_, which gives the pair counts of all feature pairs in shell *k*. **d**, log *g*_*ab*_ (*r*), pair counts over the random-label reference, of CEACAM5–CD3E on the whole section (line) with its 95% interval from the tile block bootstrap (gray band) over 500 µm tiles (Methods). Shading, level 0 (linear bins below 4 µm) and alternate lattice levels with four linear bins per log-hierarchical level. **e**, The same pair on serial Xenium and Visium HD sections of P2 and on Xenium for P1 (2 µm bins from the intra-bin shell, 0–2 µm, plotted at 1 µm).

Every value carries a standard deviation from a block bootstrap over 500 µm tiles, evaluated in closed form while the pair counts are accumulated, and a 95% interval (Methods). For replicate groups, the tile variance is combined with the between-section variance (Methods). The output of one run is, for every pair of features and every scale bin, log *g* and its 95% tile-bootstrap interval (Methods).

### 2.2 Two-point function patterns agree across platforms and differ between patients

We first applied the estimator to colorectal cancer (CRC) serial sections that were profiled on three platforms [11]: Xenium (molecule coordinates, 422-gene panel), Visium HD (2 µm bins, whole transcriptome), and, for patient P2, Visium v2 (55 µm spots), restricted to the 408 genes common to all three. The sections are serial and were not registered, so each comparison is whole section against whole section (Methods). For three marker pairs whose cell types are co-expressed, in contact, or segregated, the Xenium and Visium HD functions agreed within their intervals from a pair-dependent scale up to the millimeter scale, and each platform resolved structure down to its own resolution (Fig. 2a).

**Figure 2.**
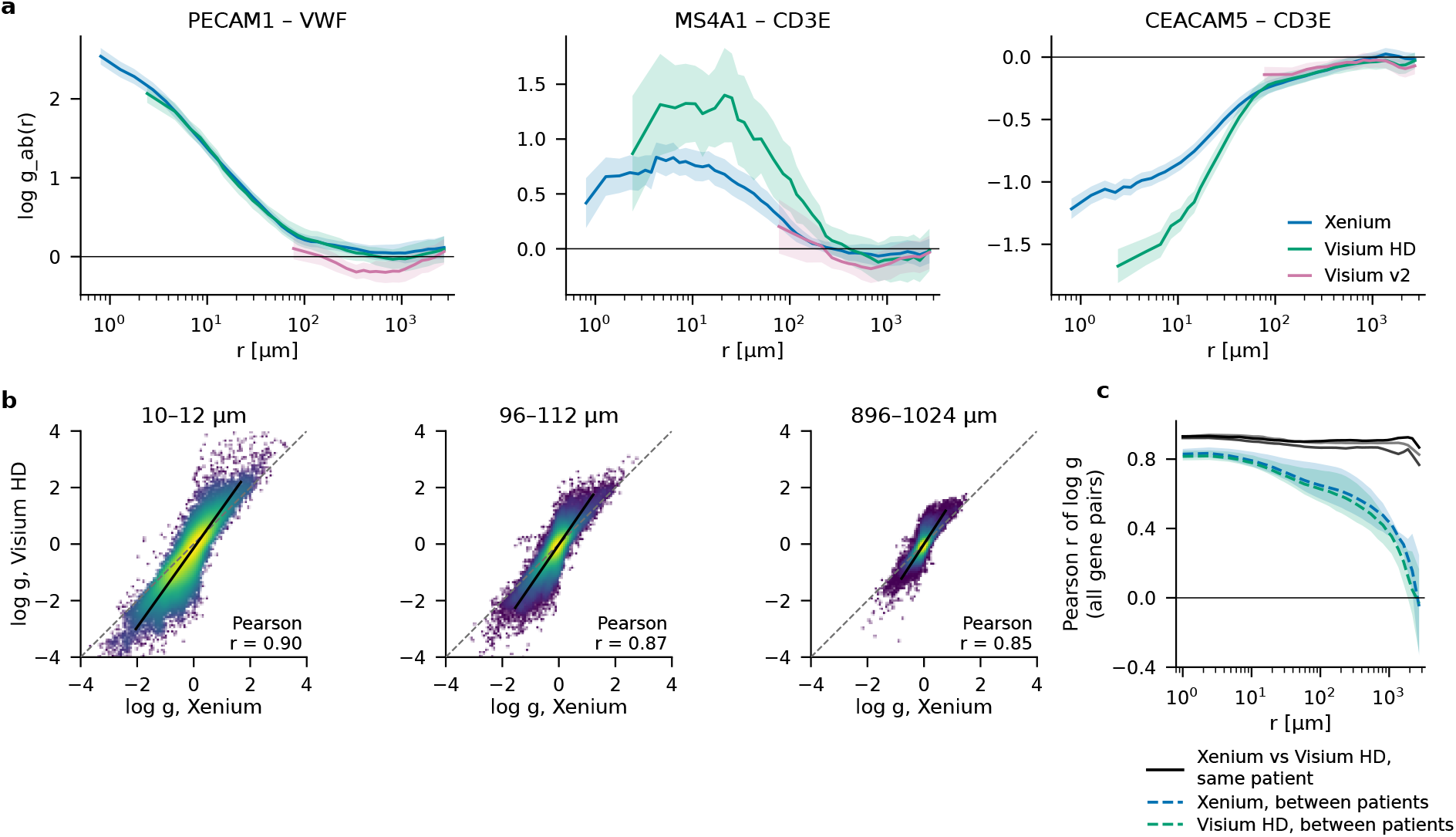
The patterns of normalized two-point functions agree across platforms on the same patient and differ between patients. Serial, unregistered CRC sections (patients P1, P2, and P5 [11]) profiled with Xenium (422-gene panel, molecule coordinates), Visium HD (2 µm bins, whole transcriptome), and, for P2, Visium v2 (55 µm spots); 408 genes common to the three platforms; bands, 95% intervals of the closed-form tile bootstrap (Methods). **a**, log *g*_*ab*_ (*r*) (random-label reference, P2) of three marker pairs of known cell-type relations, one per type of relation (two endothelial markers, PECAM1–VWF, same cells; B versus T cells, MS4A1–CD3E, contact zones; tumor epithelium versus T cells, CEACAM5–CD3E, segregated); Xenium at molecule level. Each curve starts at its platform’s resolution (Methods) and ends at 3 mm, half the section. **b**, log *g* of all gene pairs, Visium HD against Xenium, at *r* = 10–12, 96–112, and 896–1,024 µm (same axes; dashed, identity; solid, line fit; color, local density of the points, logarithmic scale; Methods); 70,341, 81,716, and 83,433 pairs with an interval and an SD below 0.5 on both platforms (44 and 9 points outside the axes at 10 and 100 µm). **c**, Replicates: Pearson correlation of log *g* over all gene pairs between Xenium and Visium HD on the same patient (solid, one line per patient) against the correlation of each platform between patients (dashed, mean of the three patient pairs; band, their range); every bin of the 2 µm lattice from the intra-bin shell, 0–2 µm, plotted at 1 µm.

Over all 70,000–83,000 gene pairs of P2 (Fig. 2b), the Pearson correlation between Xenium and Visium HD was 0.85–0.90 from 10 µm to 1 mm (Methods). The points did not scatter around the identity line, however: Visium HD spread log *g* about 1.5-fold wider than Xenium at every scale, and Visium v2 spots lay between the two (Supplementary Fig. S1).

Below about 10 µm, the platforms differed in a way that their resolution does not explain. The two endothelial markers agreed down to the smallest Visium HD bin, whereas the two pairs of markers of different cell types deviated (Fig. 2a), and CEACAM5–CD3E was more segregated on Visium HD, the coarser platform, than on Xenium (Fig. 1e); a loss of resolution would instead flatten log *g*. This residual, as well as the substructure of the point clouds in Fig. 2b, could arise from a platform-dependent, cell-type-specific efficiency of detection or permeabilization that acts at and below the cell scale (cell-type permeabilization, or “CP”, symmetry breaking). Such an effect would be a property of the platforms rather than of the tissue of the two serial sections.

The platform agreement held on all three patients and exceeded the agreement between patients (Fig. 2c). Xenium and Visium HD agreed with Pearson *r* = 0.91–0.94 from 1 to 6 µm on every patient and 0.84–0.91 from 0.1 to 1 mm, whereas the same platform on different patients correlated with 0.79–0.87 from 1 to 6 µm and 0.25–0.77 from 0.1 to 1 mm. The between-patient standard deviation of log *g* exceeded the within-section bootstrap error 1.8–3.3-fold at *r* ≤ 256 µm, so the patient differences were resolved far above the measurement error.

A second tissue set showed where this agreement ends: on mouse spleen sections profiled with Xenium, Visium HD, and MERSCOPE [56], cross-platform agreement over 4,095 gene pairs on the same animal was lower than the agreement of one platform between two infected animals (Supplementary Fig. S2; Methods). Below the 2 µm bin pitch, the Xenium–Visium HD correlation dropped only in the intra-bin shell on the spleen (0.71 against 0.78 at 6 µm) and not at all on the CRC sections, so the comparison needs no lower distance cutoff. Cross-platform agreement is therefore a quantitative statement, to be compared with the biological contrast of interest.

### 2.3 Molecule-level functions resolve a knockout effect that cell-level analyses miss

We next asked whether molecule-level functions resolve a spatial change that cell-level analyses miss, on nine Xenium mouse spleen sections from the SpatialBench study [56]: three wild-type (WT), three *Tbx21*^fl/fl^ *Cd23*^cre^ knockout (KO), in which the transcription factor T-bet is deleted in mature follicular B cells [57], and three *Cd23*^cre^ control (Ctrl; *Tbx21*^+/+^) spleens, 100-gene panel. The SpatialBench authors chose this model because loss of T-bet in B cells disrupts the polarization of germinal centers into dark and light zones, which also impacts the relative distribution of B and T cells [56, 57]. B and T operators were defined per molecule from marker genes and, for the cell-level analyses, per segmented cell with the same markers (Methods, Operators and Baselines). At cell level, we ran Squidpy co-occurrence with its recommended settings [18], and we evaluated our estimator on cell centroids with cell types as features, which is equivalent to spatstat’s cross-*K* function [37].

All three representations showed the same shape of the B–T function above ≈10 µm, but only the molecule-level function extended below the cell diameter (Fig. 3a). The molecule-level B–T anti-correlation was deeper than the centroid one on every section (log *g* = ™0.44 versus ™0.18 at 10 µm, median over nine sections; Fig. 3d), and its 95% interval was narrower, by a median factor of 2.1–2.4 at 10–50 µm (Fig. 3b).

**Figure 3.**
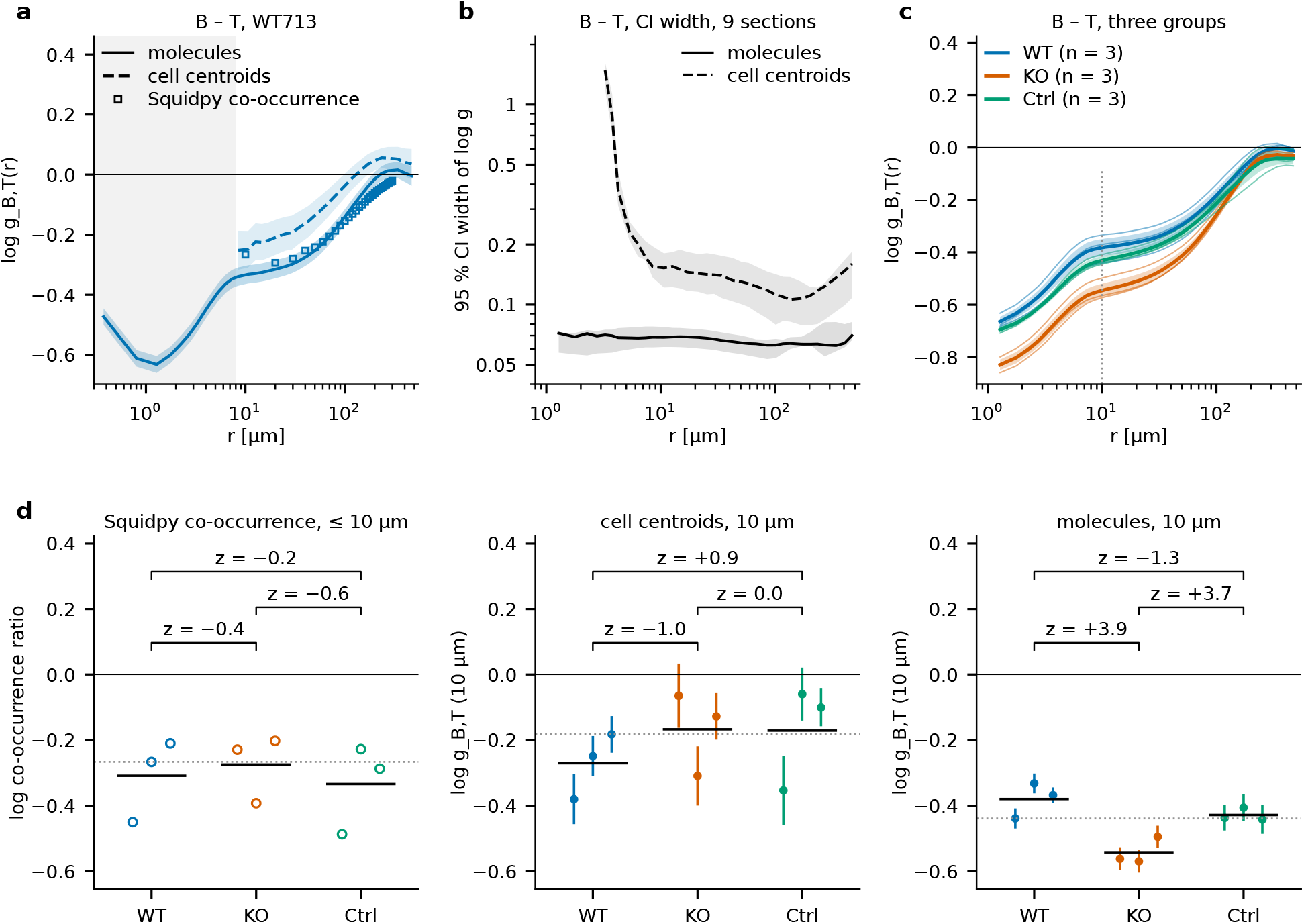
Molecule-level functions resolve structure below the cell diameter and a knockout effect missed at cell level. SpatialBench mouse spleen [56] (Xenium, 100-gene panel; three wild-type (WT), three *Tbx21*^fl/fl^ *Cd23*^cre^ knockout (KO), and three *Cd23*^cre^ control (Ctrl) sections). B and T operators are defined per molecule from marker genes (Methods) and, for the cell-level analyses, per segmented cell with the same marker sets. Line style gives the level: solid, molecules; dashed, cell centroids with the identical estimator. **a**, log *g*_B,T_(*r*) on WT713 and Squidpy `co_occurrence` on 50,000 cells (squares; ratio for the cell pairs within distance *r*); bands, 95% intervals; gray,below the cell diameter (8 µm), where the centroid function is not shown. **b**, Width of the 95% interval of log *g*_B,T_: median over the nine sections (lines) and interquartile range over sections (bands). The centroid curve extends down to the smallest distance bin in which at least five of the nine sections have an interval (3.0–3.5 µm); its width rises steeply below the cell diameter, where few cell pairs contribute. **c**, Molecule-level log *g*_B,T_(*r*) per section (thin) and group mean ± error (tile bootstrap and between-section spread; *n* = 3 per group) by group (color); dotted, 10 µm, the distance of d. **d**, B–T at 10 µm per section for Squidpy co-occurrence (log ratio as in a, cell pairs within 10 µm; open circles), the cell-centroid function, and the molecule-level function (distance bin 10–12 µm; filled circles with 95% tile-bootstrap intervals); black bars, group means; gray dotted, median over the nine sections (™0.27, ™0.18, and ™0.44). Brackets give *z*, the difference divided by its error, of WT™KO, Ctrl™KO, and Ctrl™WT; the error combines the tile bootstrap and the between-section spread (Squidpy, between-section spread only; Methods). Ctrl™WT is the calibration contrast, in which no knockout effect is expected.

The three groups provided a differential test with three sections per group and a built-in calibration (Fig. 3c,d). Ctrl and WT both carry intact *Tbx21*, so a contrast counts as a knockout effect only if WT™KO and Ctrl™KO both reach |*z*| ≥ 2 with the same sign while Ctrl™WT does not (*z*, difference divided by its error; Methods). At 10 µm, B and T molecules avoided each other more strongly in KO than in WT and Ctrl spleens (WT™KO = +0.16±0.04, *z* = 3.9; Ctrl™KO = +0.11±0.03, *z* = 3.7; Welch’s *t*-test on the three per-section values, *p* = 0.017 and 0.025), while Ctrl™WT was ™0.05 ± 0.04 (*z* = ™1.3; Fig. 3d). Neither cell-level analysis resolved this difference: the knockout contrasts of the centroid function (*z* = ™1.0 and 0.0) and of Squidpy’s co-occurrence within 10 µm (*z* = ™0.4 and ™0.6) lay within about one error of zero.

Dark and light zones of germinal centers are characterized by their composition of B (and T) cells, so a disrupted zone structure is expected to change the B–T function [57, 58]. The groups began to separate at about 100 µm (WT™KO *z* = 2.5, Ctrl™KO *z* = 1.8), the length scale of these zones, and both knockout contrasts reached |*z*| ≥ 2 at all shorter distances, while Ctrl™WT stayed below |*z*| = 2 at every scale (Fig. 3c). The difference is therefore what a disturbed zone structure predicts; it is resolved at the molecule level and missed at the cell level.

### 2.4 Tissue area and replicates, not molecule depth, set the statistical error

To predict the precision of a measurement, we subsampled tissue tiles and fitted σ(log *g*) ∝ *f* ^™ *α*^ to the error at area fraction *f* (Methods). If tiles contributed independently, *α* would be 0.5, so that four times the area would halve the error. On a CRC Xenium section, a Xenium 5K lymph node [59], and four MERFISH liver sections [60], *α* was 0.41–0.50 at every scale tested (Fig. 4a,b). The error therefore falls close to the square root of the sampled area in all tested cases.

**Figure 4.**
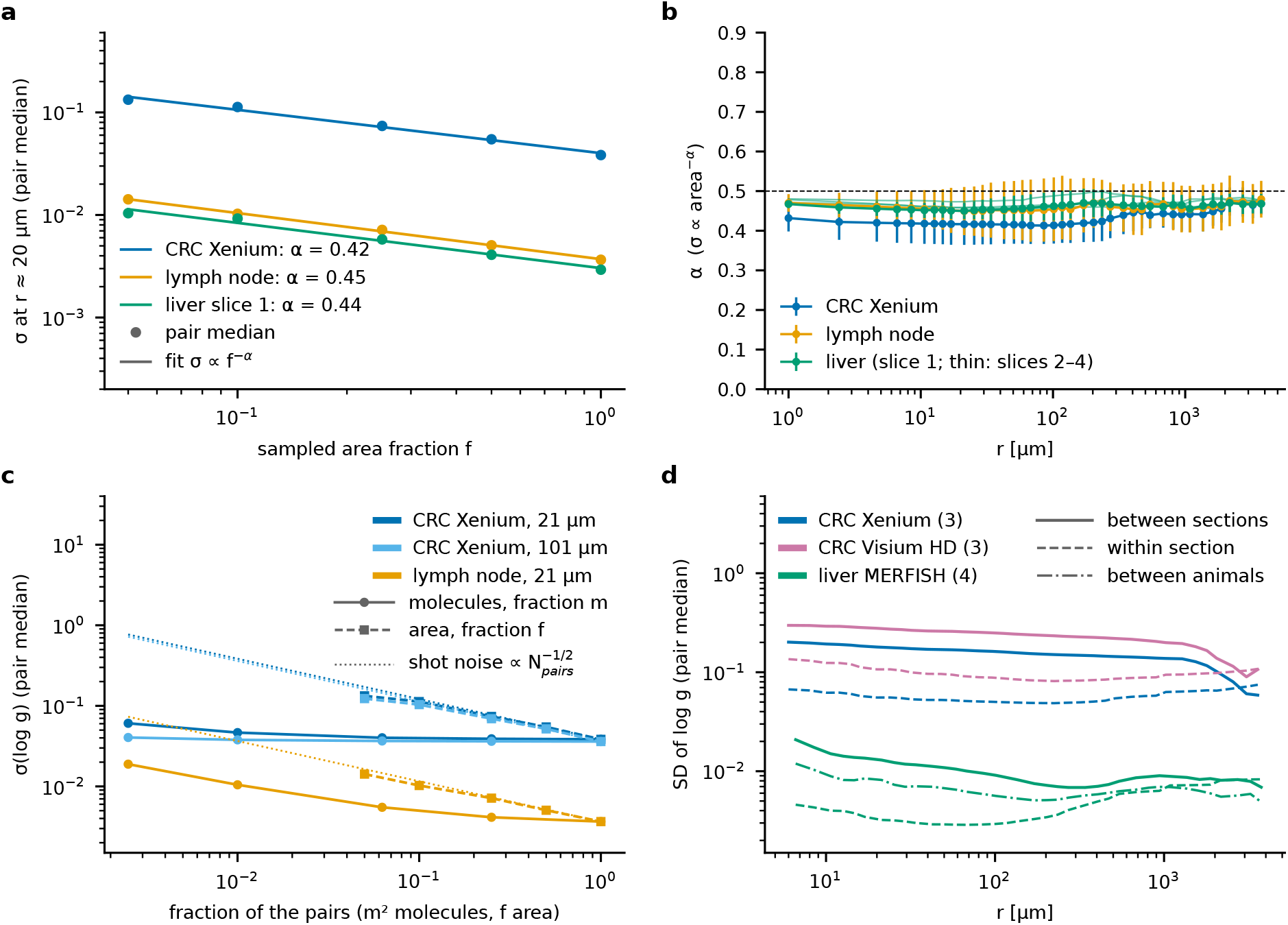
Statistical error of scale-resolved correlation functions: area and replicates, not molecule counts, set the precision. **a**, Standard deviation of log *g*_*ab*_ (*r*) at *r* ≈ 20 µm (closed-form tile bootstrap within the subset; median over the 20,100 pairs of the 200 most abundant genes on every tissue) against the sampled area fraction *f* for CRC P2 Xenium (48 mm^2^), a Xenium Prime 5K human lymph node (61 mm^2^), and MERFISH mouse liver section 1 (100 mm^2^; areas, bounding boxes of the molecules); points, pair medians; lines, least-squares fits of σ ∝ *f* ^™ *α*^, with *α* in the legend (Methods). **b**, The exponent *α* against *r* for the same data (error bars, SD over 8–16 independent tile-subset draws; thin lines, liver sections 2–4); dashed, *α* = 0.5, the value for independent tiles. **c**, Molecules versus area on the CRC section (*r* ≈ 21 and 101 µm) and the lymph node (21 µm). A fraction *m* of the molecules gives *m*^2^ of the pairs, so both routes are plotted against the fraction of the pair count: σ(log *g*) when molecules are subsampled at fixed area (solid) or area is subsampled at fixed density (dashed); dotted, pure shot noise 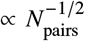. **d**, Replicates: between-section SD of log *g* (solid) against the within-section bootstrap SD (dashed) for three CRC patients on Xenium and Visium HD serial sections and for four MERFISH liver sections (two animals × two serial sections; dash-dotted, SD of the two animal means).

At fixed area, the number of molecules mattered much less (Fig. 4c). On all three tissues, the dependence on the molecule fraction was nearly flat at full depth (Methods), so deeper sampling of the same section cannot reduce the error much further. A real re-sequencing of one Visium HD library confirmed this (Supplementary Fig. S3; Methods): threefold depth lowered the error 1.7-fold at 4 µm but at most 1.11-fold at *r* ≥ 48 µm, and the same sequencing budget spent on a third of the area gave a larger error at every scale above 4 µm.

Replicates set the cohort-level error (Fig. 4d). The between-patient SD of log *g* on the three CRC patients (0.14–0.19 on Xenium) was well above the within-section error, whereas four MERFISH liver sections from two mice agreed 13–22-fold more closely, with no between-animal component resolved. For data producers, the recommendation is therefore to spend budget on tissue area and on the number of sections, and on depth only where structure below about 20 µm is the question.

### 2.5 Platform artifacts are confined to characteristic scales

At molecule level, platform artifacts occupy characteristic bands of distance instead of biasing every downstream count. We found three classes: loss of same-gene pairs below the optical resolution, genespecific backgrounds, and losses to segmentation (Fig. 5; Methods). On the CRC Xenium section, samegene pairs were missing below 1 µm, whereas different-gene pairs showed no deficit (Fig. 5a,b). The likely causes are optical crowding and the decoder’s suppression of duplicate calls (Methods), so samegene functions on Xenium are read from 1 µm on.

**Figure 5.**
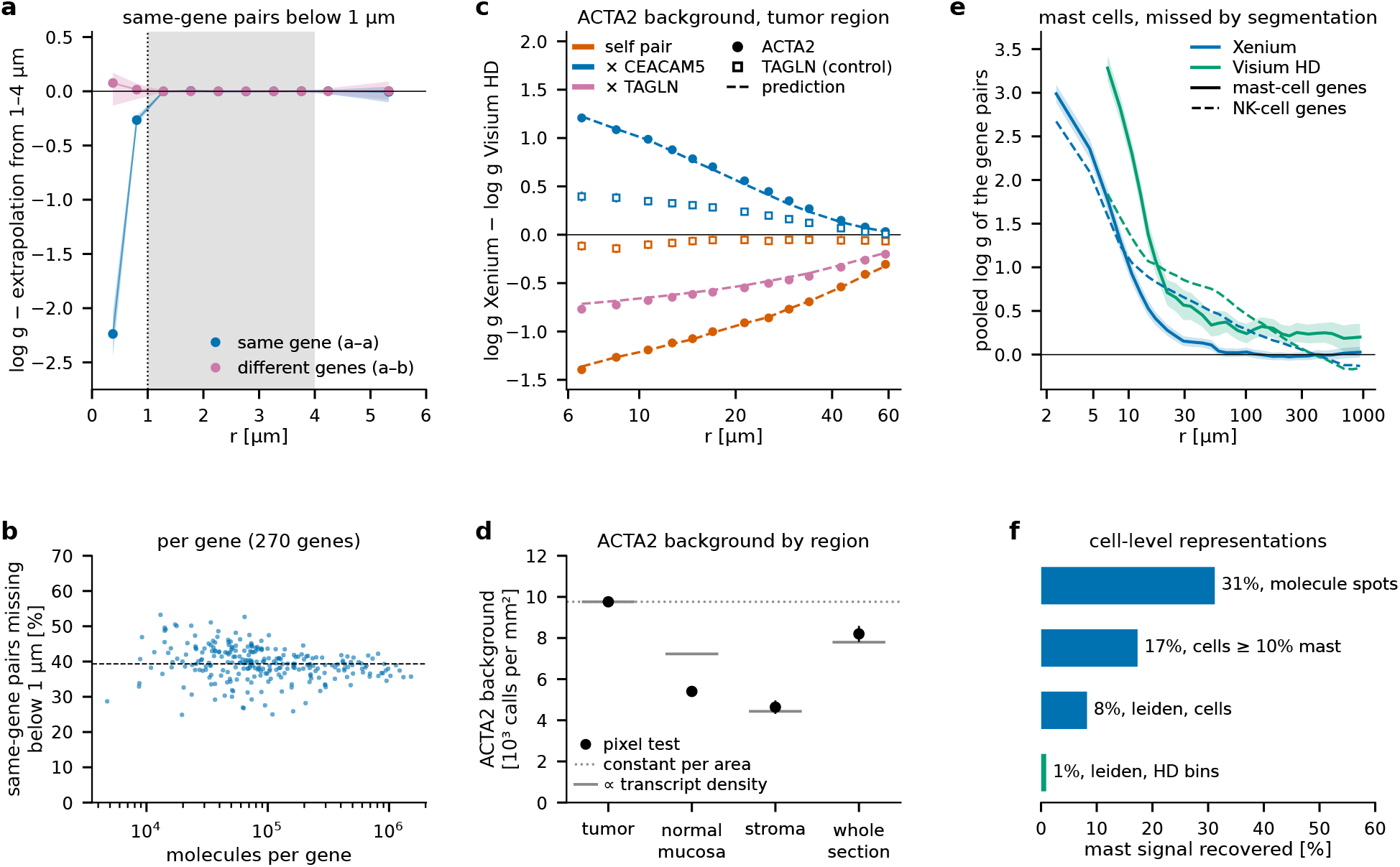
Platform artifacts are identifiable, scale-localized features of the molecule-level functions. Xenium CRC P2 [11], 408 genes, and Visium HD on the serial section (Methods). **a**, log *g* minus a quadratic extrapolation from 1–4 µm (gray) for same-gene (270 genes) and different-gene pairs (50,175), median and interquartile range over pairs; negative values mark pairs missing relative to the extrapolation; dotted line, 1 µm. **b**, Fraction of same-gene pairs missing below 1 µm per gene against its abundance (molecules per gene); dashed, median. **c**, Xenium™Visium HD difference of log *g* in the tumor region for the ACTA2 pairs (filled) and the TAGLN pairs (open, control); color,partner gene; error bars, combined tile-bootstrap SDs; dashed, prediction for the background fraction *b* measured by the pixel test. **d**, Density of the ACTA2 background from the pixel test in the tumor, normal-mucosa, and stroma regions and on the whole section (all tissue pixels); error bars, SD of the closed-form tile bootstrap; two models scaled to match the tumor value: a constant density per area (dotted) and a density proportional to the total transcript density (solid). **e**, Pooled log *g*(*r*) of the 21 pairs of seven mast-cell genes (solid) and, for comparison, of four genes of NK cells, a dispersed cell type (dashed), on Xenium (molecules in 2 µm bins) and Visium HD (2 µm bins); color, platform; bands, 95% intervals of the closed-form tile bootstrap. **f**, Share of the mast signal recovered by celllevel representations: fraction of mast molecules in dense molecule spots (DBSCAN), fraction of segmented mast transcripts in cells with at least 10% mast content, and purity of the purest cluster on segmented cells and on Visium HD 8 µm bins.

A gene-specific background was the second class. Inside the tumor region, Xenium’s ACTA2 deviated from Visium HD on all of its pairs, although its quality scores were above the panel median. A pixel-level test attributed 64.6 ± 0.9% of the Xenium ACTA2 calls to a background independent of the gene’s co-expressed markers, 2,300-fold the rate per codeword of the negative controls (Methods). Taking Visium HD as background-free, a background of that size, spread like the random-label reference, predicted the cross-platform difference of the three ACTA2 pairs at every scale from 6 to 60 µm without a free parameter (Fig. 5c). The background was not uniform per area: it followed the total transcript density in the stroma and on the whole section, as expected for calls that scale with the local RNA content, and was 25% below it in the normal mucosa (Fig. 5d; Methods). One background fraction therefore predicts the ACTA2 pairs within a region, not across the whole section.

The third class is segmentation. The 21 pairs of seven mast-cell genes all ranked in the top 1% of 83,028 gene pairs on both platforms at 2–7 µm (Fig. 5e). Cell-level representations recovered little of this signal: 83% of the segmented mast transcripts sat in cells with less than 10% mast content, and no cluster of a standard clustering pipeline recovered mast cells (Fig. 5f). For exclusive markers of a rare, compact cell type, the molecule-level function at these distances is a segmentation-free detector.

At the sub-micrometer end, three effects limit what same-gene functions measure, which we quantified on an AVITI24 (Cytos) cell-culture data set [7, 61] and a public CosMx section of human kidney [5, 62] (Supplementary Fig. S4; Methods). First, the random-label reference permutes labels over the whole sample, so differences in expression between cells score as same-gene clustering. Permuting labels only among the molecules of each segmented cell measures this composition pedestal. At 0.5–1.0 µm, on the same 184 genes, the sub-cellular excess above the pedestal was 88% of the same-gene signal on AVITI24 and 34% on CosMx (Supplementary Fig. S4a). The share is a property of the sample, not of the estimator. Second, projecting AVITI24’s seven z-planes before counting raised the same-gene log *g* in the first bin from +1.02 (per plane) to +2.18, that is, from 2.8-fold to 8.9-fold enrichment in *g* (Supplementary Fig. S4b). Third, on CosMx, no same-gene pairs lay closer than 0.10 µm, and below 0.5 µm the same-gene curve was non-monotonic, a decoding artifact, which places the usable floor at about 0.5 µm (Supplementary Fig. S4c).

### 2.6 Billion-molecule analyses on a single GPU

We benchmarked the estimator on identical real inputs against two references that accept molecules (Fig. 6; Methods): Squidpy `co_occurrence` with genes as clusters, which is designed for cell clusters and is used here outside its intended setting because no published tool computes all gene pairs on molecules, and a KD-tree cross-*K* reference, the estimator behind `spatstat Kcross` and `pointpats`. Runs used one H100 GPU or 16 threads of its host CPU; every SCOTE run returned the pair functions of all gene pairs with their uncertainties, and its time runs to the result in host memory (Methods). For the whole CRC Xenium section (47.1 million molecules) and the 200 most abundant genes over 48 shells to 16.4 mm, this took 13.5 s on the GPU and 63 s on 16 CPU threads. A whole-transcriptome Visium HD section (18,085 genes) took 14.1 min on the GPU (Fig. 6b). A single call computed all pairs of all 18,028 genes of one FFPE cervical-cancer section (10x Atera preview, 1.04 billion molecules [63]) with their uncertainties in 26.2 min, with a peak of 53.4 GB of allocated device memory. Cost followed area rather than molecule count (Fig. 6c).

**Figure 6.**
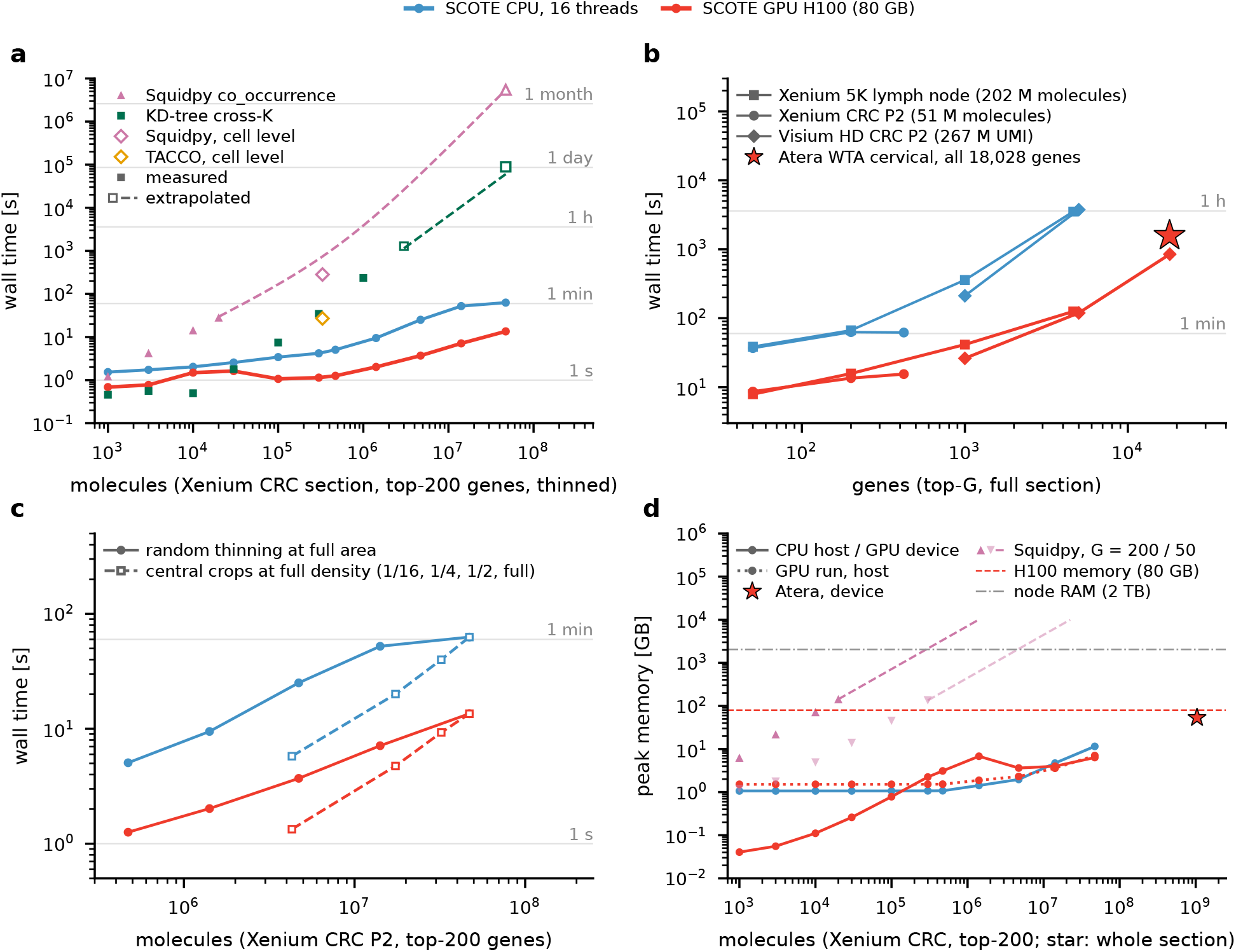
Compute scaling on real data with molecule-level comparators. Single runs without IO. One NVIDIA H100 (80 GB) with 16 host threads, or 16 pinned threads of its host CPU, an Intel Xeon Platinum 8480CL (colors, top legend, all panels); every SCOTE run returns log *g* of all gene pairs with its closed-form tile-bootstrap uncertainty. **a**, Wall time versus molecules at fixed area (Xenium CRC P2, 200 most abundant genes, uniform thinning; identical inputs for all methods); full section 13.5 s on the H100 and 63 s on 16 threads. Squidpy `co_occurrence` at molecule level (16 threads; triangles) and the KD-tree cross-*K* reference (16 workers; squares): filled, measured; open, extrapolated (KD-tree, from part of the gene pairs, full section ≈25 h from 32% of the pairs; Squidpy, full section ≈63 days); dashed, model extrapolation. Open diamonds, cell-level co-occurrence of the 331,383 segmented cells (20 clusters, to 300 µm) with Squidpy (283 s) and TACCO (27 s). **b**, Wall time versus gene-panel size on full sections: Xenium 5K lymph node (up to 202 million molecules), Xenium CRC P2 (up to 51 million), and Visium HD CRC P2 (2 µm bins, 267 million UMIs, up to 18,085 genes; whole transcriptome 14.1 min on the H100). Star, one Atera WTA FFPE cervical-cancer section with all 18,028 genes, 1.04 billion molecules in one call on the H100 (26.2 min). **c**, Area versus density: thinning at full area (solid) against central crops at full density (dashed; 1/16, 1/4, 1/2, and the full area); at equal molecule numbers the crops are 1.2–4.0-fold cheaper. **d**, Peak memory versus molecules: CPU host memory and GPU device memory (solid), host memory of the GPU runs (dotted); star, device peak of the Atera section (53.4 GB); Squidpy measured (200 and 50 genes; triangles) and its buffer (4*N LG*^2^ bytes, dashed, drawn up to 10^4^ GB); horizontal lines, H100 device memory and the 2 TB RAM of the node.

Extrapolated from runs up to 300,000 molecules, Squidpy’s co-occurrence with 16 threads on the full section would need ≈63 days and a 362 TB buffer. The KD-tree reference would need ≈25 h on 16 workers, extrapolated from 32% of the gene pairs. With uncertainties, the references were faster only below a few 10^4^ molecules (Fig. 6a). At the cell level, its intended use, Squidpy’s co-occurrence of the 331,383 segmented cells of the same section took 283 s, and TACCO [29] took 27 s. The molecule-level computation for all pairs of 200 genes with their uncertainties on the GPU therefore cost less than a celllevel analysis. SCOTE’s GPU and CPU paths give bit-identical counts, with and without uncertainties.

## 3 Discussion

Normalized two-point functions give spatial omics a common quantitative currency: their patterns transfer between platforms, resolutions, and representations, with or without a segmentation. Where platforms differed, the difference was attributable and, apart from a platform-specific amplitude of log *g*, localized in scale. Because the estimator is the same for every platform and every scale, these effects become measurable features instead of hidden biases.

Scale-resolved two-point functions thus enable differential analysis with errors and a calibration contrast; cross-platform comparison at stated normalization levels and scales; experimental design by tissue area and replicates; all gene pairs at all scales for a billion molecules and more; and quality control in the bins where artifacts act. Their output, tables of pair, scale, value, and interval, can be stored with the data [64, 65].

However, the implementation is currently two-dimensional, which biases the first bins on thick samples. While the approach can pinpoint artifacts, it cannot correct for them, so affected bins are simply discarded. Our analyses are based on cohorts of a few samples or on single samples, we took vendor quality filters and segmentations as given, and all timings are based on single runs.

In conclusion, scale-resolved two-point functions computed at molecule level provide one readout across platforms and resolutions, with errors that can be planned and artifacts that can be named. Nothing in the estimator is specific to RNA: it needs only labeled positions or binned values, so the same code applies directly to proteins and to any other molecule-resolved or binned spatial modality, which we have not yet done systematically.

## 4 Methods

### 4.1 Data sets and common data model

All data are public (Data availability). The SpatialBench spleens come from mice infected with *P. berghei* ANKA, treated with chloroquine and pyrimethamine, and harvested 12 days after infection; wild-type mice were female, knockout and control mice male [56, 66]. Every platform is converted once into one of two on-disk representations, with coordinates in µm. The *molecule table* (parquet; imaging platforms) holds `x, y, [z]`, `feature`, `sample`, `cell_id` (™1 for unassigned), `nucleus`, and `qv`. Xenium `transcripts.parquet` is filtered to molecules with qv ≥ 20 and stripped of control codewords. The 10x Atera preview transcripts (zarr store) are filtered in the same way. MERSCOPE/MERFISH `detected_transcripts.csv` provides global *x, y* and the z-slice index; blank barcodes are dropped. The *bin table* (sparse CSR counts per bin and feature, bin coordinates in µm, bin size) holds Visium HD `square_002um` outputs (8 and 16 µm bins for the mast-cell comparison) and Visium v2 filtered matrices with spot positions. Segmented cells (Xenium `cell_feature_matrix.h5` and centroids) are used for the baselines only. Gene panels are intersected before any cross-platform comparison (CRC: 408 genes common to Xenium, Visium HD, and Visium v2; spleen: 90 genes common to Xenium, MERSCOPE, and Visium HD).

### 4.2 Two-point estimator

For features *a, b*, and a distance bin *k, P*_*ab*_ (*k*) is the number of ordered pairs (*i* ∈ *a, j* ∈ *b, i* ≠ *j*) with *r*_*ij*_ in bin *k*. The normalized two-point function is

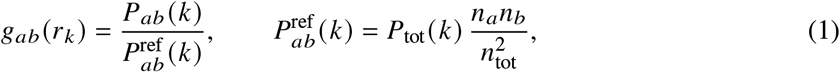

where *P*_tot_(*k*) is the pair histogram of all objects regardless of label and *n*_*a*_ the number of objects of feature *a*. This random-label reference is the expectation under a permutation of feature labels among the observed positions (random labeling), so that *g*_*ab*_ is the normalized mark connection function of the labeled pattern [36, 50, 51]; it removes geometry, edge effects, tissue holes, and density gradients at the cost of one extra histogram and is used throughout. We report log *g*_*ab*_ (*r*); the connected function is *g* ™ 1. For bin tables the objects are bins weighted by their counts (*X* is a count matrix); for molecule tables *X* is 0/1 per molecule at level 0 and a count matrix per lattice cell at levels ≥ 1.

*Distance binning*. At level 0 (imaging platforms), molecule pairs are enumerated directly in linear bins [0, Δ), …, [7Δ, 8Δ) with Δ = 0.5 µm. At levels *L* ≥ 1, molecules are binned onto a lattice of pitch *h*_*L*_ = 2Δ· 2^*L*™1^, features are aggregated per lattice cell into a sparse count matrix *X*_*L*_, and lattice-cell pairs are counted by their integer lattice distance in the four shells [4*h*_*L*_, 8*h*_*L*_), which continue seamlessly from the previous level’s maximum; the next level halves the integer coordinates. With 12 levels this gives 8 + 4 · 12 = 56 bins from 0 to 16.4 mm (four bins per octave). For binned platforms and for imaging data binned to a common grid (Figs. 2b,c, 4, 5e) the hierarchy is lattice-only: level 0 consists of shells 0–7 at the bin pitch (2 µm; shell 0 holds intra-bin pairs, *X*^⊤^ *X* minus the diagonal), followed by 11 coarse levels (52 bins, *r* < 32.8 mm; bins beyond the section size are empty). The discretization bias of lattice shells relative to annuli (up to ±17%) cancels exactly in *g*, because numerator and reference are evaluated on the same lattice. In Fig. 2a, each curve starts at the first bin beyond its platform’s resolution: beyond the 0.5 µm localization bin for Xenium, beyond the 2 µm bin for Visium HD, and at the first neighbor shell of the 64 µm spot lattice for Visium v2.

### 4.3 Shell products on the log-hierarchical lattice

At level *L* the pair histogram of all feature pairs in shell *k* is the matrix product

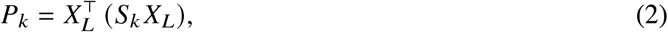

with *S*_*k*_ the sparse (cells × cells) shell adjacency (1 where two occupied lattice cells are at a lattice distance in shell *k*). The cost is therefore linear in the number of molecules at fine levels, where *X*_*L*_ is sparse, and that of a dense matrix multiplication at coarse levels. *S*_*k*_ is built from the sorted hashes of the occupied lattice cells. Shells are lattice shells, *k* = ⌊|*o*|⌋ of the integer cell offset *o* at each level. Because the offset at level *L* + 1 is half that at level *L* only up to the parities of the two cells, the windows of consecutive levels overlap, and the shell counts sum to ≈2–6% more than the number of pairs, depending on the pair-distance spectrum of the section (5.1% for the CRC Xenium section), rather than partitioning them. Each shell’s count is exact for its window, and *g* is a ratio within one window whose random-label reference uses the same window, so the overlap cancels exactly in the normalization. Fits across shells use the joint tile-bootstrap covariance (see below) rather than treating shells as independent. The product is evaluated by one of three back-ends, chosen per level by the density of *X*_*L*_: (i) two sparse–sparse products (SciPy; the reference implementation [67]); (ii) a fused parallel kernel (numba [68]) with per-thread *G* × *G* accumulators and BLAS matrix multiplication above 5% density (used for all CPU per-tile runs); (iii) on the GPU (PyTorch [69]), fine levels as row-blocked sparse–dense products (nnz(*X*) · *G* multiply-adds per shell) and coarse levels as dense matrix multiplication, with an out-of-memory split fallback and fp64 accumulation. All paths are exact: the counts are integers below 2^53^, so fp64 products are bit-identical to the sparse reference (verified in the test suite and on the Visium HD and Xenium CRC sections). The output is one *G* × *G* slab per shell, accumulated in fp64 on the device. Its per-tile parts feed the accumulators of the tile bootstrap (see below); for the gene subsets and operators of the analyses here, which relabel molecules before counting (see below), they are also kept as a tile tensor (the per-tile pair counts, tiles × bins × *G* × *G*) for statistics across pairs and shells. For whole transcriptomes no tile tensor is formed, and the output is streamed shell by shell into memory-mapped files, so only one shell’s accumulators are in memory at a time.

### 4.4 Block bootstrap over tiles

Every section is cut into square tiles of 500 µm (81–356 tiles per section). All quantities of the estimator are sums over tiles: the pair count 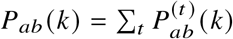, the all-label histogram *P*_tot_(*k*), and the feature totals *n*_*a*_ and *n* = *n*_tot_. Each pair is split in half between the tiles of its two members (symmetrized per-tile slabs 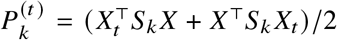, with *X*_*t*_ the rows of *X* in tile *t*), so that the SD of log *g*_*ab*_ equals that of log *g*_*ba*_; point estimates do not depend on this attribution. A block bootstrap draws *T* tiles with replacement *B* times and recomputes log *g*, including its reference, from the resampled sums, so both features are resampled jointly by tile [52, 53, 55, 70]. We evaluate this bootstrap in closed form.

For a tile-additive quantity *x* = ∑_*t*_ *x*_*t*_, the variance over multinomial resampling of the tiles is Var(*x*) = ∑_*t*_ (*x*_*t*_ ™*x*/*T*)^2^, its value for infinitely many resamples. Linearizing log *g*_*ab*_ = log *P*_*ab*_ ™log *P*_tot_ ™log *n*_*a*_ ™ log *n*_*b*_ + 2 log *n* gives the contribution of tile *t*,

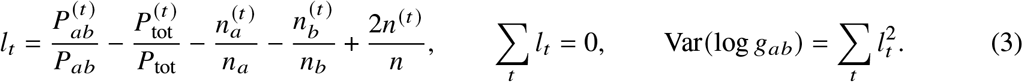

This closed-form (linearized, *B* → ∞) tile block bootstrap is an infinitesimal jackknife over tiles [70]. Because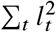 expands into sums over tiles of the per-tile slab times weights that are known before the pass, it is accumulated in four *G* × *G* arrays per shell while the pairs are counted, and no tile tensor has to be stored. The SD is free of resampling noise but not exact: as an estimate from a finite number of tiles, the variance has a relative error of order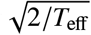, and like every tile bootstrap it ignores correlations between neighboring tiles. We report log *g* from the full-section counts with the central 95% interval log *g*±1.96 SD. Its reliability is set by the effective number of tiles that carry the pair, *T*_eff_ = *P*^2^/∑_*t*_ (*P*^(*t*)^)^2^: for *T*_eff_ ≥ 20 the interval is reported; for 5 ≤ *T*_eff_ < 20 it is reported and flagged; for *T*_eff_ < 5 no interval is given. Pairs without an interval are left out of every pair set used for statistics across pairs, including the SD filters. In the convention of a Monte Carlo bootstrap that attributes each pair to the tile of its first member, the median ratio of the closed-form SD to a 200-resample bootstrap is 1.003 for *T*_eff_ ≥ 20 (Xenium CRC P2), and the closed-form SD is 2.5–5% smaller for 5 ≤ *T*_eff_ < 20. Resampling bootstraps were run only as checks of the closed form: on this whole section, and on subsets of 10–20 tiles (see below), where a few tiles carry most of a pair’s counts and the closed-form SD is 8–24% below a 2,000-resample bootstrap. Statistics over many pairs (Pearson’s *r*, Deming and OLS slopes, RMS and mean differences, pooled pair sets) and ratios of tile sums are smooth functions of tile sums; their SD follows from the same per-tile contributions by the delta method, Var(*θ*) = ∑_*t*_ (∑_*ab*_ (∂*θ*/∂ log *g*_*ab*_) *l*_*t*_ (*ab*))^2^ with the pair set fixed, summed over independently tiled sections. Fits across shells (the background fits of Fig. 5) use generalized least squares with the cross-shell covariance of the tile bootstrap, Cov(*k, k*^′^) = ∑_*t*_ *l*_*t*_ (*k*) *l*_*t*_ (*k*^′^), with its correlation matrix shrunk toward the identity (Ledoit–Wolf), because neighboring shells correlate at 0.9–1.0 and the smallest eigenvalues of a covariance estimated from the tiles of one or two sections are biased low.

### 4.5 Normalization and region operators

The feature totals in the reference remove a uniform per-gene capture efficiency, but not differences in resolution, cell-type-dependent detection efficiency, segmentation, or section content. Cross-platform comparisons are made between whole sections at the same scale (global), and restricted to the same compartments only where stated (the artifact tests, Fig. 5c,d). Compartments are defined by one platform-independent region operator: marker-set fractions of the common genes on a 10 µm grid, smoothed with a Gaussian of σ = 50 µm, divided by the smoothed total count, z-scored over tissue pixels, argmax over regions, applied identically to molecules and bins. CRC regions: tumor, normal mucosa, and stroma, each defined by a marker set.

### 4.6 Cross-platform agreement metrics (Fig. 2b,c and Supplementary Figs. S1 and S2)

*Agreement* (Fig. 2b,c): per bin, over the pairs *a* ≤ *b* with an interval (*T*_eff_ ≥ 5) and a tile-bootstrap SD below 0.5 on both platforms, the Deming slope of log *g*_*B*_ on log *g*_*A*_ with error-variance ratio *λ* = median 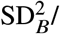median 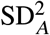(tile-bootstrap SDs), and Pearson *r*; intervals *θ* ± 1.96 SD by the delta method over the per-tile contributions (see above), with the pair set and *λ* fixed and the two sections independent. Points are colored by their local density (binned Gaussian kernel density estimate); the line fit is drawn where the 95% interval of its slope excludes zero. The Visium v2 spots (55 µm diameter, 100 µm center to center on a hexagonal array) are counted on a square analysis lattice of 64 µm pitch (2^5^×2 µm, so that its shells coincide with the 2 µm hierarchy from 256 µm on). Its first neighbor shell, 64–128 µm, contains only nearest-neighbor spot pairs and is compared with the sum of the four 2 µm-lattice bins in that range.

### 4.7 Sequencing-depth ablation (Supplementary Fig. S3)

For the depth ablation, the deep 2 µm matrix was binomially thinned to 0.335 (the UMI total of the minimum-depth run), 0.5, 0.25, and 0.125 (UMI totals and reads in Supplementary Fig. S3). All depths share the section and its bins, so the tiles are identical at every depth; on the 79 tissue tiles of the section the error was computed as described below for the 200 most abundant genes, at full area for every depth and on area subsets of the deep run.

### 4.8 Composition pedestal, z-projection, and resolution (Supplementary Fig. S4)

The *within-cell permutation* permutes feature labels among the molecules of the same segmented cell (incell molecules only, per z-plane, three permutations, seed 0) and evaluates the same estimator; its log *g* is the composition pedestal, the observed minus the pedestal is the sub-cellular excess, and the share is excess/observed. CosMx data: GEO GSE282059 [62] (human kidney, CosMx 6k panel), tissue piece 2 of the seven on the slide (fields of view 132–198; 53.8 million endogenous molecules, 119,027 cells). The cross-platform pedestal comparison used the 184 genes that are among the 1,000 most abundant genes of both data sets (mitochondrial genes and control probes excluded). *Stacked* same-gene functions are count-weighted averages over genes, 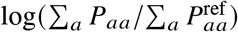. *Per-plane versus projected*: pairs are counted only within the same z-plane, or after projecting all planes. *Resolution*: the sub-micrometer curve was computed in 0.05 µm bins on the first 16 fields of view of the piece.

### 4.9 Operators (spleen, Fig. 3)

Operators label molecules: B = Cd19, Ighd, Ighm; T = Cd3e, Cd3d, Trac, Cd8a. The operator run is the all-pairs two-point run on the relabeled molecule table (pair enumeration at Δ = 0.5 µm plus 8 lattice levels, *r* ≤ 1 mm, 500 µm tiles).

### 4.10 Baselines (Fig. 3)

Both cell-level analyses run on the same nine sections with the same marker sets, on the vendor segmentation (97,000–360,000 cells per section). *Cell typing*: *f*_*k*_ = counts of marker set *k* divided by the panel counts of the cell; B or T by the larger of *f*_B_ and *f*_T_ (at least 0.1), for cells not first labeled as follicular dendritic cells (FDC), T follicular helper cells (Tfh), or germinal-center B cells from their marker fractions; unlabeled cells are kept as “other” so that the reference uses all cells. *Squidpy 1*.*8*.*3* with its recommended imaging settings: `co_occurrence` with 31 distance edges from 0 to 300 µm on a random subsample of 50,000 cells (the routine is quadratic). The ratio reported at edge *r* counts all cell pairs within distance *r*, so the curve is cumulative; Fig. 3d uses the value for the pairs within 10 µm. *Centroid-level g*_*ab*_ (*r*):the identical two-point pipeline on cell centroids with feature = cell type; not interpretable below the cell diameter (≈8 µm), where few cell pairs contribute.

### 4.11 Replicate groups (Fig. 3c,d)

For groups of sections (Fig. 3c,d), the group value is the mean of the per-section log *g*, with error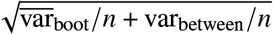 for *n* sections per group, where 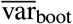 is the mean closed-form tile-bootstrap variance of the sections (see above); contrasts are quoted as difference ± error and *z* = difference/error. Squidpy’s co-occurrence has no per-section error, so its error is the between-section term alone.

### 4.12 Error scaling with area, molecules, and replicates (Fig. 4)

*Area*: from a section’s tile tensor and for *f* ∈ {0.05, 0.1, 0.25, 0.5, 1}, round(*f T*) tiles are drawn without replacement, σ_*f*_ (*r, ab*) is the closed-form tile-bootstrap SD of log *g*_*ab*_ (*r*) within the subset (see above; on the tensor restricted to the 200 genes, so the reference is that of the panel), averaged over 8 (single pairs: 16) independent subset draws, log σ_*f*_ = ™*α* log *f* + *c* is fitted per pair, and the median *α* over the 20,100 pairs of the 200 most abundant genes is reported per bin; its error is the SD over subset draws. Two terms bias the within-subset SD low: the factor 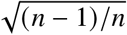 a bootstrap over *n* tiles, which lowers *α* by 0.01– 0.02, and, where a few tiles carry most of a pair’s counts, the linearization shortfall of the closed-form SD (see above). On the CRC section, the within-subset *α* falls up to 0.09 short of 0.5, mostly through these two terms. Bins where shells leave the section (*r* ≳ 4–5 mm) are not quoted. *Molecules*: the run is repeated on seeded uniform subsamples *m* ∈ {0.05, 0.1, 0.25, 0.5} at full area (reference recomputed).

The area route at equal pair count is read from the *f* -subsets; it was 1.7–1.9-fold noisier than the molecule route. *Replicates*: for CRC P1, P2, and P5, the between-patient SD σ_obs_ (median over about 83,000 pairs) contains the within-section noise 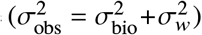 and is compared with the within-section bootstrap SD σ_*w*_. For the four MERFISH liver sections (two animals, two serial sections each), the between-section SD (*v* = 3), the pooled within-animal SD (*v* = 2), and the SD of the two animal means (*v* = 1) are compared with the bootstrap SD.

### 4.13 Artifact tests (Fig. 5)

*Sub-resolution collapse*: from the molecule-level CRC P2 Xenium run (0.5 µm bins), log *g* of every samegene pair (genes with at least 20 pairs in each bin up to 4 µm) and every different-gene pair is fitted over 1–4 µm (bins 2–7) by a quadratic in *r* and extrapolated to 0–0.5 and 0.5–1 µm; the residuals *d*_0_, *d*_1_ are the deficits (median and IQR over pairs), the missing-pair fraction of a bin is 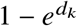, and below 1 µm it is 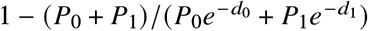 with the observed pair counts *P*_*k*_. The threshold is set at 1 µm, two bins. *Gene-specific background*: inventory of the raw transcript table including control codewords, quality scores, and in-cell fractions per gene; the pixel intercept test bins each domain (the tumor, normal-mucosa, and stroma regions, and the whole section, that is, all tissue pixels regardless of region) to 10 µm pixels, keeps the pixels with at least one gene call, and divides the mean ACTA2 count in pixels without TAGLN or DES calls by the mean over the kept pixels, which estimates the co-marker-independent fraction *b* of the domain (SD from the closed-form tile bootstrap of this ratio of tile sums over 500 µm tiles); the background density is *b* times the ACTA2 density of the domain. The background rate relative to the negative controls (2,300-fold) is the mean ACTA2 count in the pixels without co-marker calls (0.99 per pixel) divided by the mean count per codeword of the 61 negative-control probes and codewords in all occupied tumor pixels (4.2 × 10^™4^ per pixel). A background making a fraction *b* of gene *a*’s molecules and spread like the random-label reference (proportional to the local density of all molecules) transforms the normalized functions exactly as

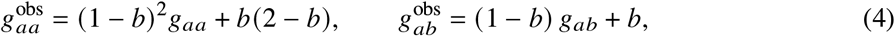

so the Xenium™Visium HD difference inside the region is predicted from *b* and the Visium HD functions (taken as background-free) and compared with the observed difference from 6 to 60 µm. *Control*: the Xenium™Visium HD difference of TAGLN–CEACAM5 is not zero (+0.40 at 6.6 µm) because strongly anti-correlated pairs differ between the platforms panel-wide (median +0.41 at the same Visium HD value); this shared part also enters the ACTA2 cross pairs (ACTA2–CEACAM5: +1.21 observed, +1.22 predicted). TAGLN is the control because it is expressed by the same smooth-muscle cells and its tumor-to-stroma density ratio agrees between the platforms (log ratio ™1.95 on Xenium, ™1.96 on Visium HD; ACTA2 ™0.73 and ™1.43); the pixel test is not informative for TAGLN, whose co-marker DES is absent from 94% of the tumor pixels. *Regions and whole section*: the ACTA2 background density from the pixel test is 9,760 ± 140 calls per mm^2^ in the tumor, 5,400 ± 170 in the normal mucosa, 4,640 ± 330 in the stroma, and 8,200 ± 390 on the whole section. The same comparison was made on the normal-mucosa and stroma regions and on the whole section (2 µm-lattice runs of both platforms), with two background models anchored on the tumor density *β* = *b x*_tumor_: constant per area, or proportional to the total Xenium transcript density. Against a constant density per area (9,760 everywhere), the measured densities are 0.55, 0.48, and 0.84 of the prediction in the normal mucosa, the stroma, and on the whole section; against a density proportional to the total transcript density (7,230, 4,430, and 7,800) they are 0.75, 1.05, and 1.05. *Rare cell type*: pooled log *g* of the 21 cross pairs of seven mast-cell genes and their ranks among all 83,028 pairs with at least 10 pairs per bin, on the 2 µm tile tensors of Xenium and Visium HD (P1 and P5 as replicates); cell level from the Xenium cell matrix (cells with at least *k* mast transcripts and at least 10% mast content); a standard scanpy pipeline [71] (cells with at least 10 counts, `normalize_total`, `log1p`, PCA 30, kNN 15, leiden [72] at resolutions 0.5–8, seed 0) scored by purity and recall against the rule “at least three mast transcripts”; Visium HD 8 µm bins likewise; DBSCAN [73] on the mast molecule positions (eps 5 µm, at least three molecules). The four bars of Fig. 5f are the fraction of mast molecules in DBSCAN spots, the fraction of segmented mast transcripts in cells with at least 10% mast content (the other 83 ± 1% are spread over 34,000 cells), and the purity of the purest leiden cluster on cells (8%, recall 2.7%) and on Visium HD 8 µm bins (1%). NK-cell genes: CCL5, KLRD1, GNLY, NKG7; mast-cell genes: CPA3, KIT, MS4A2, CTSG, HPGDS, IL1RL1, GATA2.

### 4.14 Compute benchmark (Fig. 6)

All methods received identical inputs held in memory: the molecules (or Visium HD bins) of the top-*G* genes of a section, thinned uniformly at full area (without replacement, seed 0) or cropped at full density to a central rectangle holding 1/16, 1/4, or 1/2 of the bounding-box area. SCOTE was timed from the coordinates and gene codes, including the binning onto the 2 µm lattice, to the pair functions of all gene pairs in 48 shells from 0 to 16.4 mm (8 fine shells and 10 coarse levels) in host memory. The measured time includes the copies of the result from the device to host memory. CPU runs used the fused numba kernel at sparse levels and BLAS at dense levels in fp64 on 16 threads (OpenMP, numba, MKL, and OpenBLAS thread counts set per run), each run in an allocation of 16 cores of the node. CPU runs with uncertainties set `OMP_WAIT_POLICY=PASSIVE` (default spin-waiting of the two thread pools made them 3–6 times slower). GPU runs used PyTorch with fp64 accumulation, shell operators built on the device, and 16 host threads. Data loading, numba JIT compilation, and CUDA context creation were excluded from the timings. Each result was then written to memory-mapped files; writing to RAM-backed storage took 265.6 s for the 390.1 GB Atera result. CPU and GPU counts were bit-identical in every run pair of the benchmark. Squidpy 1.8.3 `co_occurrence` was run at molecule level with gene as a categorical cluster key and the same 49 shell edges as cumulative thresholds; runs whose predicted buffer (4*N LG*^2^ bytes for *N* points, *L* thresholds, and *G* genes) exceeded 200 GB were skipped. Its full-section cost was extrapolated from a joint fit of a buffer term ∝ *NG*^2^ and a pair-loop term ∝ *N*^2^ to all measured points. The KD-tree reference counts, for every unordered gene pair, neighbors between the two molecule sets at the same edges with SciPy `cKDTree.count_neighbors`, parallel over pairs in 16 worker processes. Runs with a time budget processed the gene pairs in random order and were extrapolated linearly in the number of pairs (full CRC section: 6,502 of 20,100 pairs in 8.0 h on 16 worker processes of an Intel Xeon Platinum 8480CL, ≈25 h extrapolated). Its dependence on *N* was fitted by a power law to the points with *N* ≥ 3 × 10^4^ (*t* ∝ *N*^1.44^ on 16 workers, 200 genes). The cell-level runs used the 331,383 cells with at least 10 counts at their centroids, 20 *k*-means clusters on 30 SVD components of log-normalized counts, and distances to 300 µm on 8 threads (TACCO 0.5.0 `co_occurrence` in 10 µm steps without permutations; Squidpy with 31 edges). Wall time of the compute step and peak resident memory were recorded per run, and the GPU’s peak allocated memory was read from PyTorch. Every number is a single run (one process per measurement, hardware as in Fig. 6) and is quoted without an interval.

### 4.15 Software

SCOTE was run with Python 3.12.14, NumPy 2.5.3 [74], SciPy 1.18.1, numba 0.68.0, and PyTorch 2.14.1 with CUDA 13.0; the analyses used pandas 3.0.5, pyarrow 25.0.1, scanpy 1.12.4, anndata 0.13.3, Squidpy 1.8.3, TACCO 0.5.0, scikit-learn 1.9.1 [75], leidenalg 0.12.0, matplotlib 3.11.1, and Snakemake 9.26.1 [76] in the same environment.

### 4.16 Use of AI

Large language models by Anthropic (Claude Opus 5, Opus 5.5, Sonnet 5.5, and Fable 5.1) were used as coding and writing assistants, in particular for the further development of the initially human-written code for SCOTE, analyses, and reproducible workflows.

## Data availability

All data sets are publicly available (the Vizgen liver data after registration, and the Element Biosciences data after a request form); none were generated for this study.

- CRC patients P1, P2, P5 (Xenium, Visium HD; Visium v2 for P2) [11]: 10x Genomics data sets (https://www.10xgenomics.com/datasets); GEO GSE280318.
- SpatialBench mouse spleen (Xenium WT709, WT710, WT713, KO166, KO167, KO168, Ctrl172, Ctrl173, Ctrl174; Visium HD WT709, KO168; MERSCOPE WT709) [56]: EMBL-EBI BioStudies S-BSST2361 (doi:10.6019/S-BSST2361).
- Xenium Prime 5K human lymph node [59]: 10x Genomics data sets.
- 10x Atera whole-transcriptome preview, FFPE human cervical cancer [63]: 10x Genomics data sets.
- Fresh-frozen human ovarian cancer section (Visium HD probe-based at two depths) [77]: 10x Genomics data sets.
- Vizgen MERFISH Mouse Liver Map, Liver1Slice1/2 and Liver2Slice1/2 [60]: available from Vizgen after registration at the Vizgen data release page (https://info.vizgen.com/mouse-liver-access); cited as “Vizgen MERFISH Mouse Liver Map January 2022”.
- Element Biosciences AVITI24 (Cytos workflow) A549 TNF*α* CRISPR screen [61]: Element Biosciences data set page (https://www.elementbiosciences.com/resources/dataset-explore-multimodal-pooled-screening-on-aviti24), which offers the download after a request form.
- Human kidney (FFPE), CosMx 6k panel [62]: GEO GSE282059.

## Code availability

The SCOTE source code will be released on GitHub under the BSD 3-Clause License at https://github.com/MagesLab/scote.

## Author contributions

M.I., Y.S., and S.W.M. developed the software and carried out the data analysis; S.W.M. conceptualized and supervised the project and wrote the manuscript; all authors reviewed and approved the manuscript.

## Acknowledgements

The authors thank Karl-Peter Hopfner, Johanna Klughammer, and Gregor Witte for providing and maintaining local GPU compute resources. The authors gratefully acknowledge LMU Klinikum for providing computing resources on their Clinical Open Research Engine (CORE). The work was supported by LMUexcellent, funded by the Federal Ministry of Education and Research (BMBF) and the Free State of Bavaria under the Excellence Strategy of the Federal Government and the Länder.

## Supplementary information

**Figure S1.**
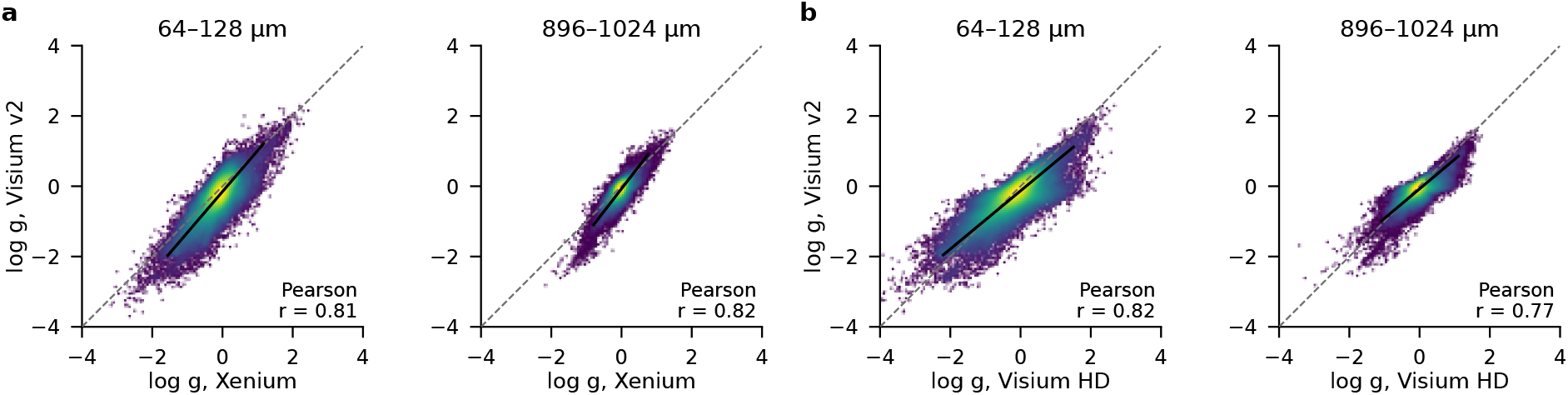
Visium v2 spots against Xenium and Visium HD, all gene pairs (CRC P2). log *g* of all gene pairs, Visium v2 (55 µm spots, 100 µm spot center distance, evaluated on a 64 µm lattice) against Xenium (**a**) and against Visium HD (**b**), at the first neighbor shell of that lattice (*r* = 64–128 µm, nearest-neighbor spot pairs only; the four 2 µm-lattice bins of the other platform in this range summed; Methods) and at 896–1,024 µm, where the spot shells coincide with the 2 µm hierarchy. Axes, identity line (dashed), line fit (solid), color (local density of the points) as in Fig. 2b. Pairs: 82,430 and 83,434 (**a**), 79,835 and 83,431 (**b**). Fitted slope of Visium v2: 1.17 and 1.29 on Xenium, 0.83 and 0.84 on Visium HD.

**Figure S2.**
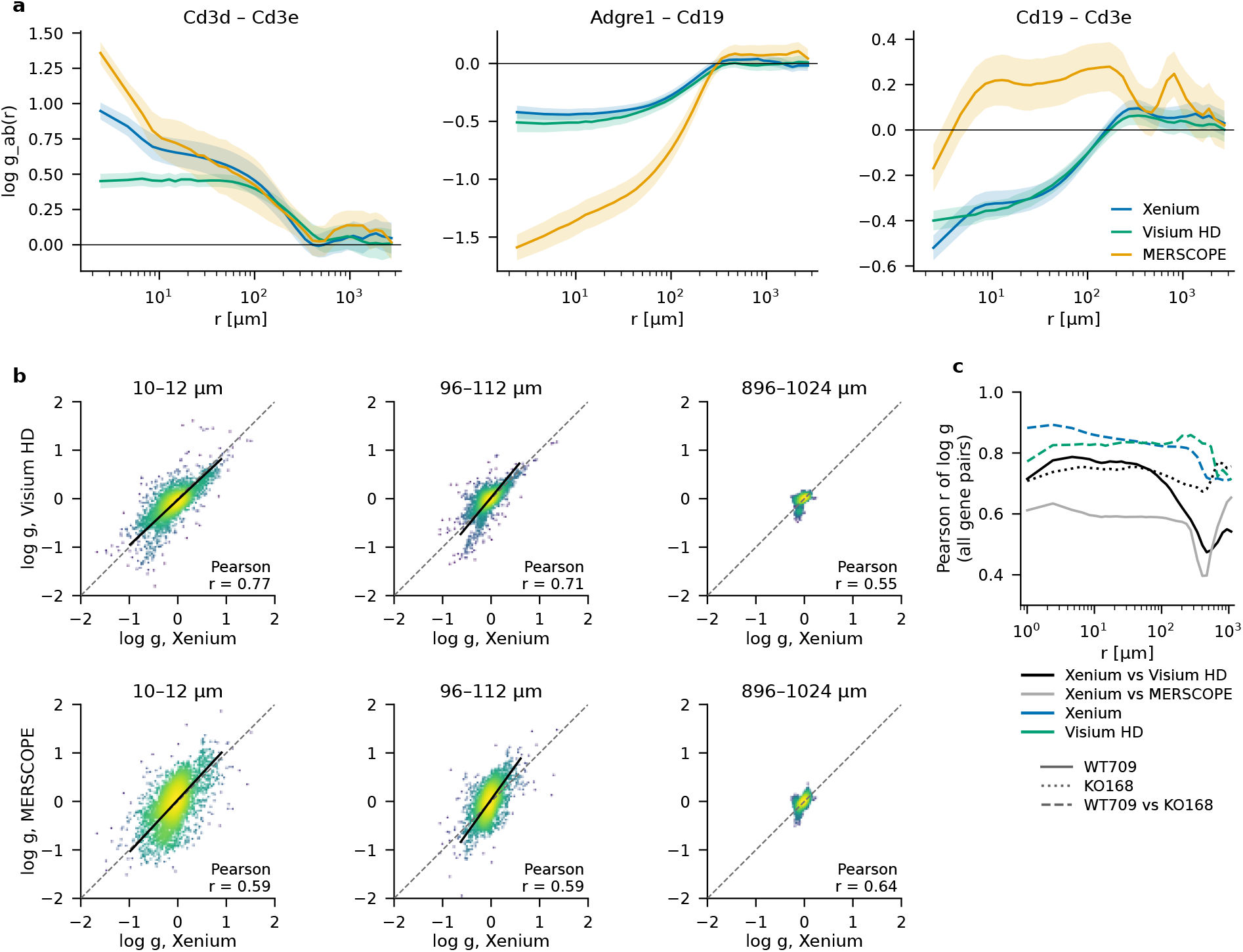
On a second tissue, cross-platform agreement falls below the agreement between two animals (SpatialBench mouse spleen, three platforms). Xenium, Visium HD, and MERSCOPE sections of the same WT709 spleen, and Xenium and Visium HD sections of a *Tbx21*-knockout spleen (KO168); 90 common genes (4,095 gene pairs); all platforms on the same 2 µm lattice; bands, 95% intervals of the closed-form tile bootstrap (Methods). **a**, log *g*_*ab*_ (*r*) of three gene pairs (T-cell co-expression Cd3d–Cd3e, red-pulp macrophage versus B Adgre1–Cd19, B–T zones Cd19–Cd3e), from the 2 µm lattice pitch to 3 mm. **b**, log *g* of all gene pairs, Visium HD (top) and MERSCOPE (bottom) against Xenium, at *r* = 10–12, 96–112, and 896–1,024 µm (same axes in all six panels, 8 points outside in each 10 µm panel; dashed, identity; solid, line fit, not drawn at 1 mm, where the 95% interval of the slope includes zero; color, local density of the points, as in Fig. 2b; 3,999–4,095 pairs per panel). Fitted slopes at the two smaller distances: Visium HD 0.94 and 1.20, MERSCOPE 1.08 and 1.39. **c**, Cross-platform agreement per animal (Xenium versus Visium HD on WT709 and KO168, Xenium versus MERSCOPE on WT709) against the between-animal agreement of one platform (WT709 versus KO168; Xenium Pearson 0.87, 0.84, 0.82, 0.81 at 6, 24, 96, 256 µm; Visium HD 0.77–0.86); the between-animal SD exceeds the bootstrap SD 1.3–3.0-fold at 1–256 µm; every lattice bin from the intra-bin shell, as in Fig. 2c.

**Figure S3.**
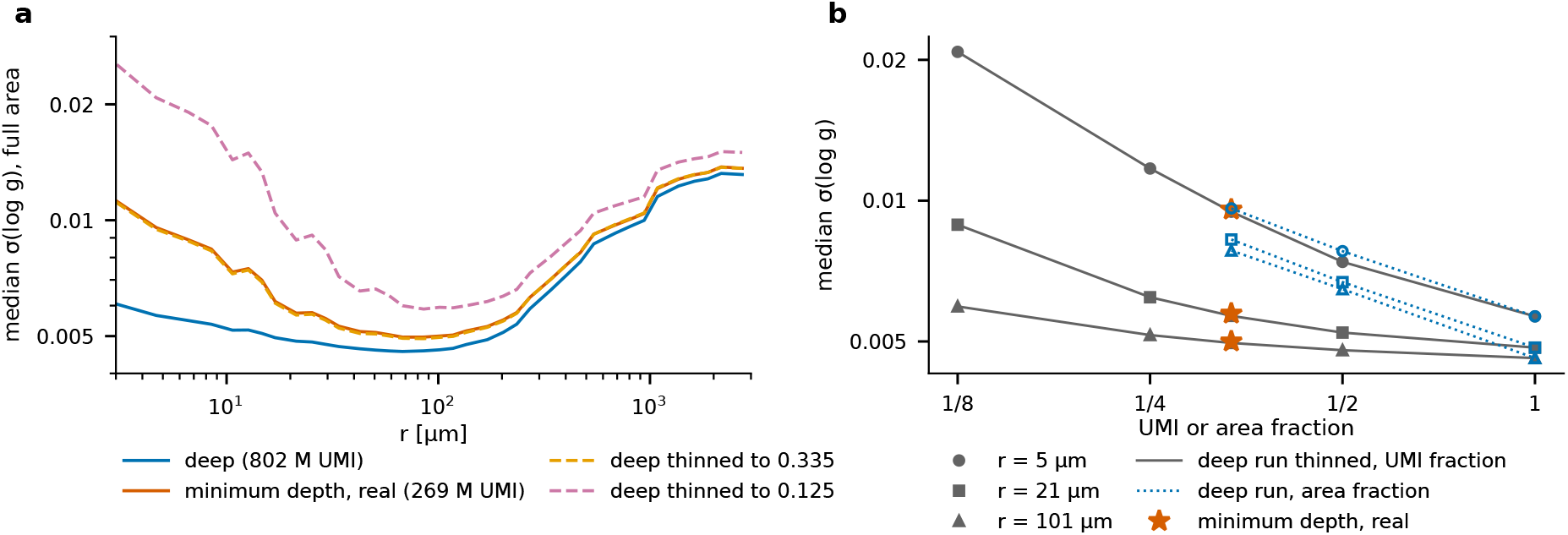
Sequencing depth buys precision mainly below about 20 µm (Visium HD probe library of one fresh-frozen ovarian cancer section). The same library sequenced to 2.32 billion reads (802 million unique molecular identifiers, UMIs; saturation 0.56) and to 0.44 billion reads (269 million UMIs, saturation 0.24), and the deep matrix binomially thinned to 0.335 (the minimum-depth UMI total), 0.5, 0.25, and 0.125. All depths share the section and its 2 µm bins, so every tile holds the same tissue at every depth and no registration is involved (79 tissue tiles of 500 µm; 200 most abundant genes; Methods). **a**, Pair-median bootstrap SD of log *g* versus *r* at full area for the deep run, the real minimum-depth run, and the deep run thinned to 0.335 and 0.125 (the real minimum-depth run and the deep run thinned to the same UMI total coincide, ratio 1.00–1.01 at every *r*). **b**, SD versus UMI or area fraction at 5, 21, and 101 µm (solid, thinned deep run; stars, real minimum-depth run) and, dotted, the same UMI budget spent on a fraction of the area at full depth.

**Figure S4.**
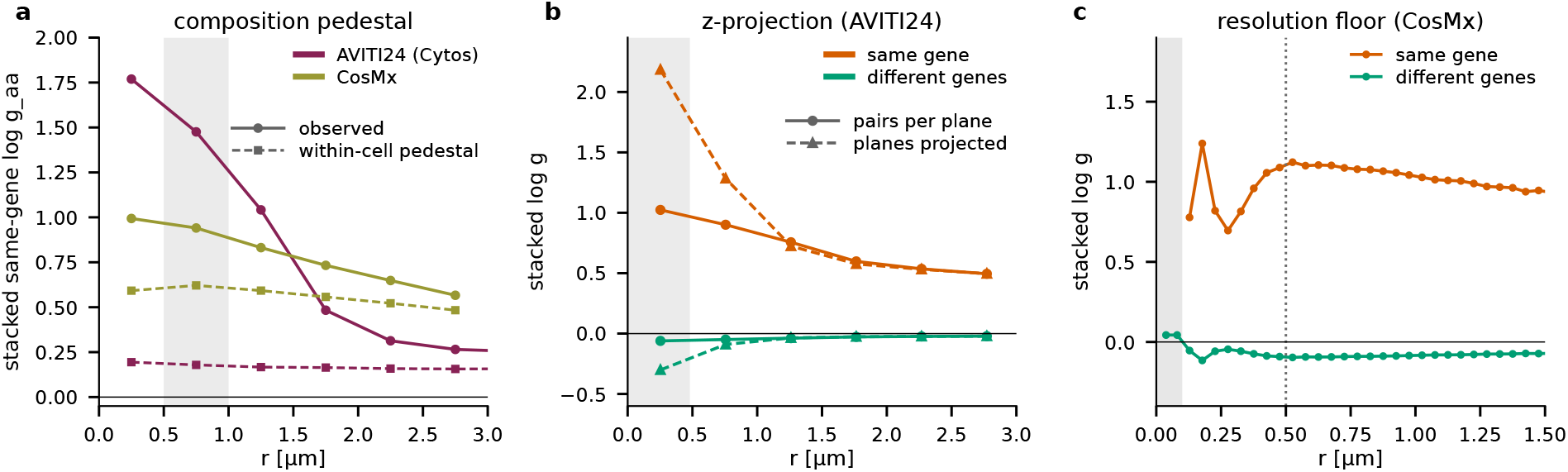
What a two-point function measures at the sub-micrometer end: composition pedestal, z-projection, and resolution floor. **a**, Composition pedestal on the 184 genes that are among the 1,000 most abundant genes of both data sets, the AVITI24 (Cytos) untargeted 3^′^ transcriptome and the CosMx 6k panel; abundant genes are used because their same-gene functions are populated below 1 µm, and the same genes on both platforms because the stacked function depends on the gene set (AVITI24 A549 cells: 46.7 million in-cell molecules in 258,419 cells; CosMx kidney: 5.35 million in 118,706 cells; in-cell molecules, pairs counted per z-plane, 0.5 µm bins on both platforms). Solid, stacked same-gene log *g*_*aa*_ against the random-label reference; dashed, the same estimator after permuting labels among the molecules of each cell (composition pedestal; mean of three permutations, whose SD, 0.001–0.010, is smaller than the line width); shaded, read point 0.5–1.0 µm. **b**, AVITI24 (50 most abundant genes, six wells, seven z-planes 1 µm apart): stacked log *g* of same-gene and different-gene pairs counted within each z-plane (solid) or after projecting the planes (dashed); shaded, offsets of at most one 0.48 µm pixel. **c**, CosMx kidney (150 most abundant genes, 16 fields of view, 0.05 µm bins): stacked same-gene and different-gene log *g* below 1.5 µm; shaded, separations without same-gene pairs (below 0.10 µm); same-gene peak +1.24 at 0.18 µm and minimum +0.70 at 0.28 µm; dotted line, usable floor at 0.50 µm.

